# Validating gene-phenotype associations using relationships in the UMLS

**DOI:** 10.1101/2020.07.29.226993

**Authors:** Andrew L Blumenfeld, Claudia Gonzaga-Jauregui, Deepika Sharma, Geisinger-Regeneron DiscovEHR Collaboration, Regeneron Genetics Center, Ashish Yadav, Shareef Khalid, Suganthi Balasubramanian, Jeffrey G Reid, Lukas Habegger, Michael N Cantor, Jeffrey Staples

## Abstract

**Objective:** Large scale next-generation sequencing of population cohorts paired with patients’ electronic health records (EHR) provides an excellent resource for the study of gene-disease associations. To validate those associations, researchers often consult databases that identify relationships between genes of interest and relevant disease phenotypes, which we refer to as simply “phenotypes”. However, most of these databases contain phenotypes that are not suited for automated analysis of EHR data, which often captured these phenotypes in the form of International Classification of Diseases (ICD) codes. There is a need for a resource that comprehensively provides gene-phenotype mappings in a format that can be used to evaluate phenotypes from EHR.

**Methods:** We built a directed graph database of genes, medical concepts and ICD codes based on a subset of the National Library of Medicine’s Unified Medical Language System (UMLS) and other resources. To obtain associations between genes and ICD codes, we traversed the defined relationships from gene, variant and disease concepts to ICD codes, resulting in a set of mappings that link specific genes and variants to these ICD codes.

**Results:** Our method created 249,764 mappings between genes and ICD codes, including 27,226 “disease” phenotypes and 222,538 “symptom” phenotypes, and provided mappings for 4,456 unique genes. Paths were validated by manual review of a diverse sample of paths. In a cohort of 92,455 samples, we used these mappings to validate gene-phenotype associations in 32,786 samples where a person had a potentially disease-causing genetic mutation and at least one corresponding diagnosis in their EHR.

**Conclusion:** The concepts and relationships in the UMLS can be used to generate gene-ICD phenotype mappings that are not explicit in the source vocabularies. We were able use these mappings to validate gene-disease associations in a large cohort of sequenced exomes paired with EHR.

## 1. BACKGROUND

When analyzing population-level cohorts that pair genetic findings with clinical data from a patient’s EHR, there is a need for a resource that provides known clinical phenotypes associated with variants in specific genes[1]. While many sources of gene-disease associations are available, they all suffer from one or more limitation that prevent them from being widely used in the validation of clinical (EHR-based) gene-disease findings. Resources like ClinVar[2], the Human Gene Mutation Database (HGMD) [3], Online Mendelian Inheritance in Man (OMIM)[4] and Orphanet[5] provide lists of diseases that have been linked to known genetic variants, either at the gene or variant level. These disease listings are a valuable reference, but the descriptions of diseases in these databases cannot be directly used to search electronic health records, where diseases are captured using billing codes such as ICD-9-CM and ICD-10-CM. Previous work has used the UMLS to elucidate general gene-disease relationships through connections involving the Gene and Human Phenotype Ontologies, but the results of these approaches are not specific to ICD codes and are therefore not specific enough to be used for validation of clinical findings[6-9]. What is needed is a way to bridge the gap between a disease name in a genetics-focused database such as OMIM and the diagnosis codes frequently used in EHRs.

In some cases, a disease name listed in a genetic database will directly match an ICD code description, but frequently the disease name does not directly correspond to an ICD code. These issues are particularly challenging in rare diseases, where ICD codes may not exist or, more often, may not be entered into the EHR with a sufficient level of granularity to distinguish among different diseases[10]. A manual approach to mapping diseases to ICD codes for thousands of genes would be very time-consuming, and a text-similarity approach would miss diseases where the appropriate ICD code description is unlike the disease name. The methodology we developed used structured medical ontologies and vocabularies to map from a genetic condition to a disease that maps to an ICD code by traversing the relationships between concepts. We believed that the relationships in these existing databases could be extracted and used to map between genes and ICD codes without significant additional manual intervention.

The need for this type of mapping can be seen in the work by Gonzaga-Jauregui, et al.[11]. The authors manually reviewed hundreds of electronic health records from the DiscovEHR [1] cohort—currently comprised of 145,000 patients from the Geisinger Health System whose exomes have been sequenced and linked to their EHR data—to identify clinical diagnoses of lipodystrophy and related diseases known to be caused by mutations in specific genes. A prerequisite for this type of analysis is a set of known gene-disease-ICD code mappings that can be used to validate the gene-phenotype associations found in large cohorts of patients such as the DiscovEHR. The analysis provided evidence that lipodystrophy is an underestimated and underdiagnosed condition in the clinical setting by identifying clinical evidence of related disorders associated with the same genetic markers. Figure 1 contains pedigrees from two families with carriers of expected pathogenetic mutations in genes that are associated with lipodystrophy and related diseases.

**Figure 1.**
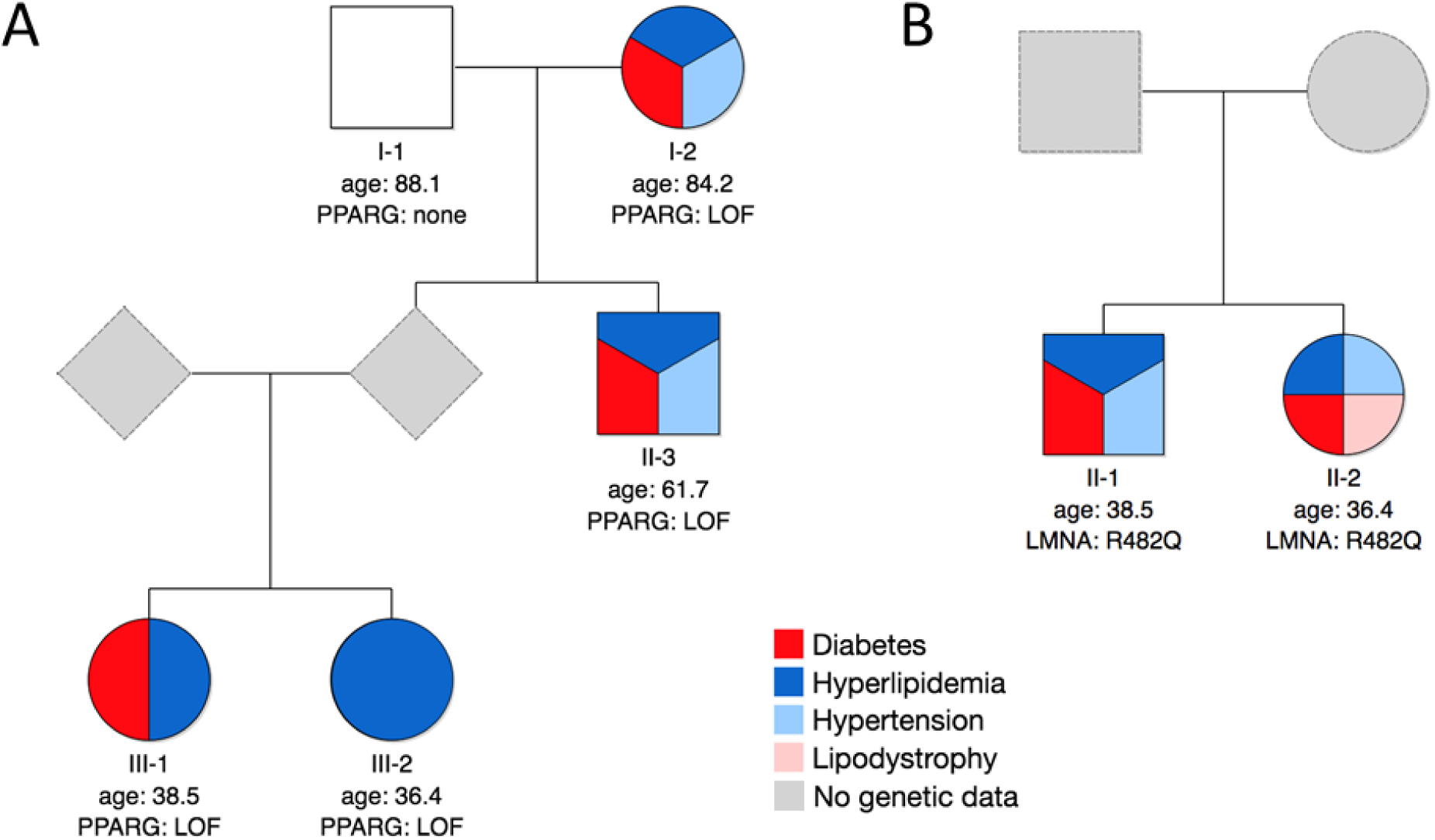
Phenotypes in carriers of predicted pathogenic variants for lipodystrophy in two DiscovEHR cohort pedigrees. The pathogenic mutations in these genes are known to result in the diseases listed in the legend. A) All carriers of the PPARG (p.E73X) mutation have one or more diagnosis codes in their EHR that corresponded to the diseases in the legend. B) Both siblings in the 2nd generation are carriers of the LMNA (p.R482Q) mutation and have diagnosis codes reported in their EHR corresponding to diabetes, hyperlipidemia, and hypertension. While the sister (II-2) is diagnosed with lipodystrophy, the brother (II-1) is not. This lack of diagnosis in the brother is not surprising as lipodystrophy is known to be underdiagnosed in males. The first line below each individual indicates the generation and the individual within that generation (i.e., II-1 indicates individual 1 of 2nd generation). Squares = males; circles = females. LOF = Loss of function variant.

Our approach leverages the relationships of existing medical vocabularies to find appropriate ICD codes for disease concepts. The Unified Medical Language System (UMLS)[12] is a compendium of medical vocabularies and ontologies that are integrated using a system of concept identifiers. Terms that are common across vocabularies are given a Concept Unique Identifier (CUI) code, and the relationships between these terms are maintained to form a network of medical concepts. By taking a subset of this network and filtering relationships for directionality we can create a path from specific, and sometimes obscure diseases to more broad disease categories and find the most appropriate ICD code. We can then use these mappings to screen a large cohort of exome variants to validate their relationship to clinical diagnoses in the associated electronic health records.

## 2. METHODS

### 2.1. Data sources

#### 2.1.1. Unified Medical Language System (UMLS)

Our primary source of data was the 2019AA release of the UMLS, and the UMLS CUIs form the backbone of our database. We downloaded and installed the MetamorphoSys program available on the UMLS website and used the files listed in Supplemental Table S1.

#### 2.1.2. MedGen

An additional source is the May 2019 release of the MedGen database. MedGen uses the same data structure and CUIs as the UMLS, although in some cases a National Center for Biotechnology Information (NCBI) CUI is used when a UMLS CUI is not available. These files can be downloaded directly from the NCBI ftp site (ftp://ftp.ncbi.nlm.nih.gov/pub/medgen/). Because MedGen uses the same data structure as UMLS, we use “UMLS” to describe the union of UMLS and MedGen data when describing the filtering processes.

#### 2.1.3. Orphanet Rare Disease ontology (ORDO)

While MedGen contains concepts from the Orphanet Rare Disease Ontology (ORDO), it does not include many of the disease-ICD-10-CM relationships available on the Orphanet web portal (https://www.orpha.net/consor/cgi-bin/Disease.php). We obtained those relationships from the ORDO ontology available at the BioPortal ontology browser[13]. To identify useful relationships, we filtered for Orphanet concepts that mapped to a UMLS CUI and an ICD-10-CM code. We only used relationship data from Orphanet-– no disease concepts from outside of UMLS and MedGen were manually added to the database.

### 2.2. Graph database

All nodes and relationships were added to a directed graph database using the GraphFrames relational graph database[14] on the Databricks Unified Analytics Platform for Genomics (a cloud computing platform built on Apache Spark) and implemented in the Scala programming language[15]. A graph database approach allowed us to accommodate the redundancy of overlapping vocabularies with the ability to add new relationships as they were identified. A directed graph database was necessary because we were interested specifically in paths from genes to ICD codes and only traversed medical concepts hierarchically from narrow to broader concepts, between synonymous concepts or from disease to symptom concepts.

### 2.3. Nodes

All nodes in our graph database represent HUGO Gene Nomenclature Committee (HGNC) gene IDs, UMLS CUIs (for genes, gene variants or medical concepts), or ICD codes (ICD-9-CM or ICD-10-CM). The relationship between these node types is depicted in Figure 2.

**Figure 2.**
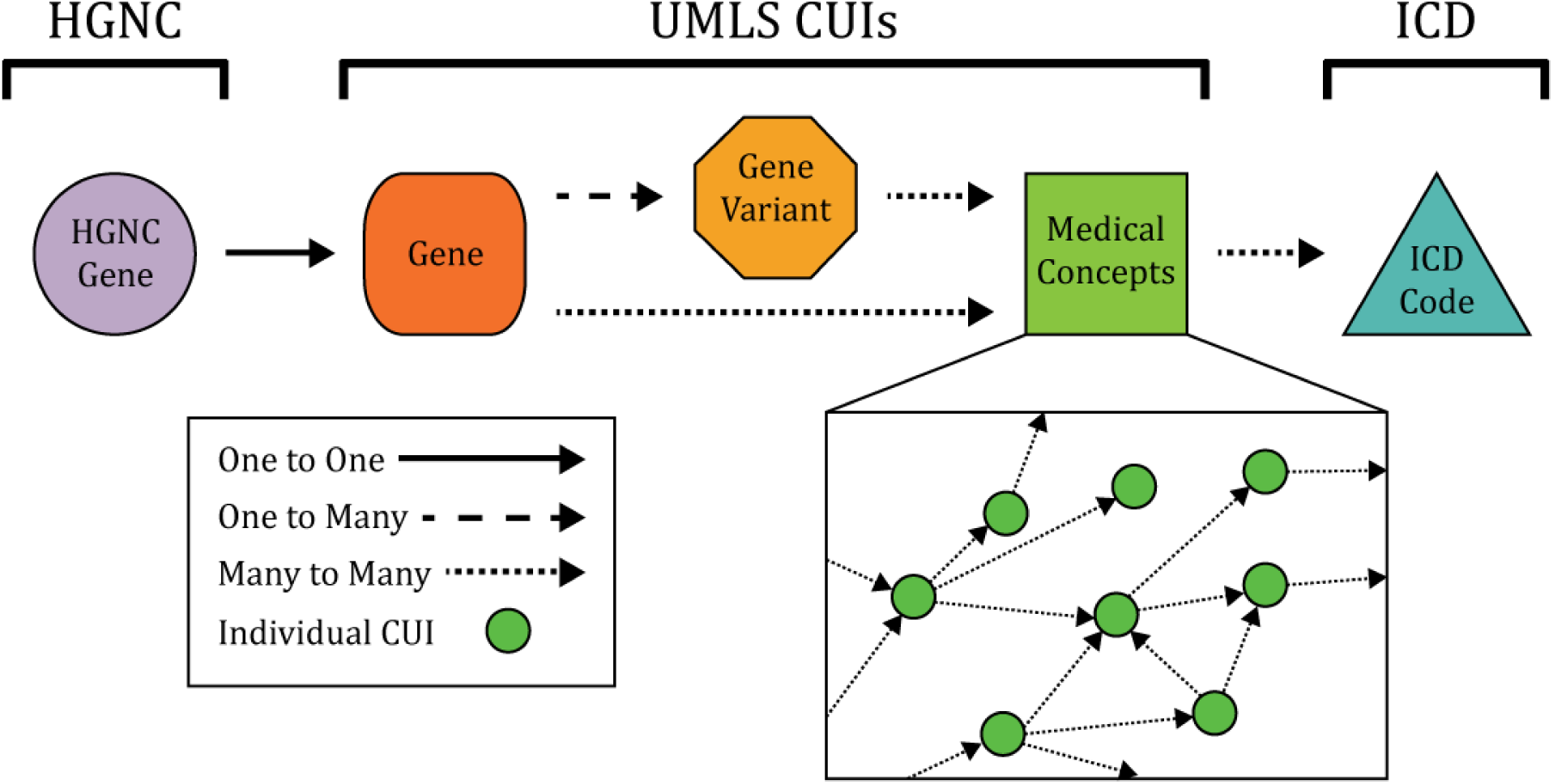
The types of nodes in the graph database and the relationships between them. All HGNC IDs are uniquely mapped to a UMLS CUI concept for that gene. Genes may map to specific variants, or directly to medical concepts. Medical concepts can be mapped to any number of ICD codes, can map to each other, and the total number of medical CUIs in a path is its “length”. The medical concepts insert shows the high-level of connectivity between the medical CUIs.

The full sets of nodes can be extracted from the UMLS using the filters in Supplemental Table S2 and the UMLS CUIs can be filtered by semantic types listed in Supplemental Table S3. Any active HGNC ID was included as an “HGNC gene” node, including protein-coding genes, pseudogenes, micro-RNAs and genomic loci[16]. For each HGNC gene node, there is a single corresponding UMLS CUI node for that gene. Additionally, many genes have known variants with their own distinct CUIs, and these variants are often mapped to diseases that are not mapped to the gene themselves. Some of these variants cannot be identified by the methods in Supplemental Table S2 and must be identified by the criteria in Supplemental Note S1.

Finally, a few nodes were labeled as “headers” because they were too broad to map to a meaningful ICD code (e.g. C0012674, “Diseases [MeSH Category]”). We were able to prevent unnecessary pathfinding and reduce the likelihood of generating incorrect paths by stopping paths at any “header” CUI (see path criteria below). Header CUIs are listed in Supplemental Table S4.

### 2.4. Edges

Edges indicate the type of relationship between CUIs and can be extracted from the UMLS files using the filters in Supplemental Table S5. Edges between HGNC gene nodes and CUIs, or CUIs and ICD codes are not broken down into subcategories. However, the UMLS contains many types of relationships between disease concept CUIs that must be filtered to a hierarchical subset[17]. In addition to hierarchical relationships between concepts, the UMLS contains relationships that can be understood as “symptoms”, which map from a disease to clinical phenotypes that are linked to that disease. The UMLS assigns a broad (“REL”) code and frequently a specific (“RELA”) code to indicate the type of relationship between the CUIs. For each combination of source, “REL” and “RELA” code, a decision was made whether to include the CUI relationships, and the full list of these decisions are in Supplemental Table S7. The “RELA” labels originate with the source vocabularies and can be in conflict between vocabularies. For this reason, there are several levels of inclusion/exclusion, and these are explained in Table 1. For example, some categories contain a mix of relationships and must be given a “soft” rejection. If some of these relationships are categorized differently by another vocabulary, they won’t be incorrectly rejected. This differs from a “hard” rejection, where the relationships in the category clearly require exclusion. For this database, we selected relationships that map a CUI to a broader CUI, to a synonymous CUI, or to a symptom CUI. Any relationships where the start and end nodes are the same and the classifications were compatible were collapsed to a single relationship, and any conflicting classifications were resolved as described in Table 2.

**Table 1:** Codes used to filter relationships

| Code | Name | Description |
| --- | --- | --- |
| G | Good | Include this relationship, indicate “conflict” if necessary |
| B | Bad (“Hard” exclusion) | Do not include this relationship, indicate “conflict” if necessary |
| E | “Soft” exclusion | Do not include this relationship, unless included elsewhere |
| Y | Include, override others | Include this relationship, ignore other sources |
| X | Exclude, override others | Do not include this relationship, ignore other sources |
| Syn | Synonym | Include this relationship, but mark as synonym (functionally equivalent to G but can be useful when evaluating results) |
| Sym | Symptom | Include this relationship, but mark as symptom |
| C | Conflict | There is a conflict between sources in how this is described |
| U | Unknown | This relationship has not been categorized |

**Table 2:** Rules for solving relationship classification conflicts

| Priority | Rules for aggregated relationship filter codes | New code description | New code |
| --- | --- | --- | --- |
| 1 | includes “X” | Do not include | B |
| 2 | includes “Y” | Include | G |
| 3 | Includes “B” and “G” | Indicate conflict | C |
| 4 | Includes “Syn” | Synonym | Syn |
| 5 | Includes "B" but NOT "G" | Do not include | B |
| 6 | Includes "G" but NOT "B" | Include | G |
| 7 | Includes "E" but NOT "G" OR "Sym" | Do not include | B |
| 8 | Includes "Sym" but NOT "G" OR "B" | Symptom | Sym |
| 9 | Other or None | Unknown | U |

#### 2.4.1. Systematically incomplete mappings

To maximize the number of successful mappings, we tried to identify relationships that were systematically lacking in UMLS. In one case, we found ∼600 CUIs where the concept contained a version of the phrase “susceptibility to”, but the concepts did not contain relationships with the root concept that they were referring to. For example, CUI C2684859 (“APLASTIC ANEMIA, SUSCEPTIBILITY TO (finding)”) was not mapped to CUI C0002874 (“Aplastic Anemia”). We were able to use the UMLS REST API to identify an appropriate CUI for over 500 of these “susceptibility to” concepts and added these relationships to the graph database.

#### 2.4.2. Other manual mappings

Approximately 150 relationships were added to the database after being identified by thorough testing. These include instances where: 1) a relationship between CUIs was missing, 2) a relationship between CUIs was incorrect and needed to be removed, or 3) a relationship from a CUI to an ICD code was missing.

### 2.5. Generating paths

The full graph database contained approximately 1.23M relationships and 666K nodes. To identify links between genes and ICD codes, we generated every possible path that fit these criteria:

- Paths must start at a node with type “HGNC”
- Paths must end at a node with type “ICD-9-CM” or “ICD-10-CM”
- Paths must have exactly one “gene” CUI node and at most one “gene variant” CUI node
- Nodes may not appear more than once in a single path
- The last CUI node in the path must be the only node that connects to an ICD node UNLESS an intermediate node connected to an ICD node and then started a symptom relationship
- The last CUI node in the path must be the only node that connects to a header CUI node UNLESS an intermediate node connected to a CUI header node and then started a symptom relationship
- A path can have at most one symptom (“Sym”) relationship
- A path can have 1 conflict (“C”) or unknown (“U”) relationship if it has a length of 1 or 2
- A path can have up to 6 CUI nodes UNLESS it has a symptom relationship, in which case it can have up to 5 CUI nodes

Any path that contained a “symptom” relationship was categorized as a “symptom path”, and all others were categorized as “disease paths”. Next, we took several steps to remove redundancy and identify a set of unique paths:

- We truncated all ICD codes to 4 digits and collapsed all similar paths to a single path
- Paths that only differed by the gene variant were collapsed into a single path
- We identified the shortest path(s) for any gene to ICD relationship (In cases where multiple paths connected the same gene and ICD code we found that the shortest paths typically included the clearest, most logical relationships between CUIs.)
- If a gene-ICD relationship occurred as both a disease and a symptom path at the same length, only the disease path was kept

As an example, the complete paths for gene P2RX2 can be found in Supplemental Figure S2.

### 2.6. Path validation

To evaluate our results, we examined several hundred paths and determined if the gene-ICD relationship was accurate and there were logical connections between the medical concepts in the path. “Disease” and “symptom” paths were evaluated separately to determine if they required separate evaluation criteria. Paths that contained “unknown” or “conflicted” edges were also evaluated separately, as these edges increased the likelihood of a path being erroneous. Using the success rate of each set, we calculated the 95% confidence interval for a sample proportion under a normal distribution[18] and the evaluation outcomes are shown in Figure 3.

**Figure 3.**
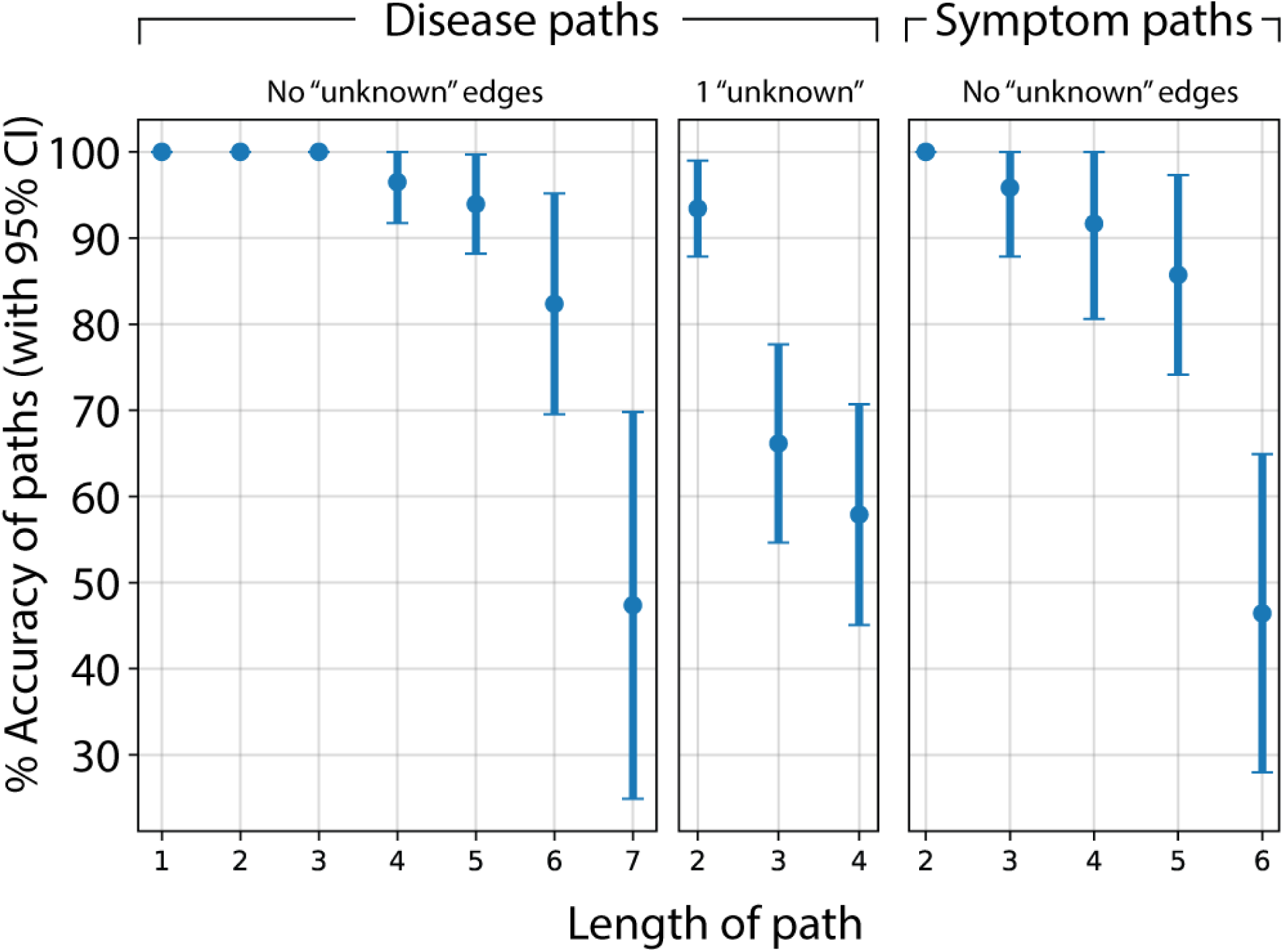
Proportion of paths that were verified as correct for each length and type, with error bars showing 95% confidence intervals. Paths were randomly sampled and evaluated for correctness of gene-ICD relationship and path logic. Disease paths with no uncertain or conflicted (i.e. “unknown”) edges show a high quality up to a path length of 6 (sample n=481). Disease paths with 1 uncertain or conflicted edge (sample n=198) only show a high quality up to a path length of 2. Symptom paths with no uncertain or conflicted edges (sample n=137) show a high quality up to a path length of 5.

When no uncertain or conflicted edges were included, we found few errors in disease paths with 3 or fewer CUI nodes or symptom paths with 2 or fewer disease CUI nodes. Our final path criteria were updated to include the following types of paths because their estimated accuracy rate was above 80%:

- Disease paths up to length 6 with no uncertain/conflicted edges
- Disease paths up to length 2 with 1 uncertain/conflicted edge
- Symptom paths up to length 5 with no uncertain/conflicted edges

## 3. RESULTS

The final counts for gene-ICD mappings are shown in Table 3. The distribution of lengths for unique mappings (same gene and 3 or 4-digit ICD code) for “disease” and “symptom” paths are shown in Figures 4A and 4B, respectively. Additionally, we evaluated the specificity of our mappings by calculating the number of unique ICD-10-CM codes that map to each gene, and vice versa. We plotted the cumulative ranks for each gene and ICD-10-CM ICD code separately for “disease” and “symptom” paths in Figure 4C-F. Figure 4C shows that in “disease” paths approximately 50% of genes map to just one ICD-10-CM code, and approximately 85% map to 10 or fewer. Figure 4E also shows that approximately 30% of ICD-10-CM codes map to a single gene. The corresponding plots for ICD-9-CM codes are included in Supplemental Figure S1.

**Table 3:** Summary of the gene-ICD paths broken down by symptom paths, disease paths, ICD-9-CM codes, and ICD-10-CM codes.

| **Description** | **Count** |
| --- | --- |
| Unique gene-ICD mappings | 249,764 |
| Unique gene-ICD disease mappings | 27,226 |
| Unique gene-ICD symptom mappings | 222,538 |
| Genes with 1 or more paths mapped | 4,456 |
| Genes with 1 or more ICD-9-CM paths mapped | 4,254 |
| Genes with 1 or more ICD-10-CM paths mapped | 4,451 |
| Genes with 1 or more disease paths | 3,772 |
| Genes with 1 or more ICD-9-CM disease paths | 2,018 |
| Genes with 1 or more ICD-10-CM disease paths | 3,758 |
| Genes with 1 or more symptom paths | 4,134 |
| Genes with 1 or more ICD-9-CM symptom paths | 4,046 |
| Genes with 1 or more ICD-10-CM symptom paths | 4,109 |
| ICD-9-CM (collapsed to 3-digit) codes mapped | 1,677 (524) |
| ICD-9-CM (collapsed to 3-digit) codes mapped with disease paths | 950 (371) |
| ICD-9-CM (collapsed to 3-digit) codes mapped with symptom paths | 1,408 (480) |
| ICD-10-CM (collapsed to 3-digit) codes mapped | 2,905 (929) |
| ICD-10-CM (collapsed to 3-digit) codes mapped with disease paths | 1,748 (642) |
| ICD-10-CM (collapsed to 3-digit) codes mapped with symptom paths | 2,345 (852) |

**Figure 4.**
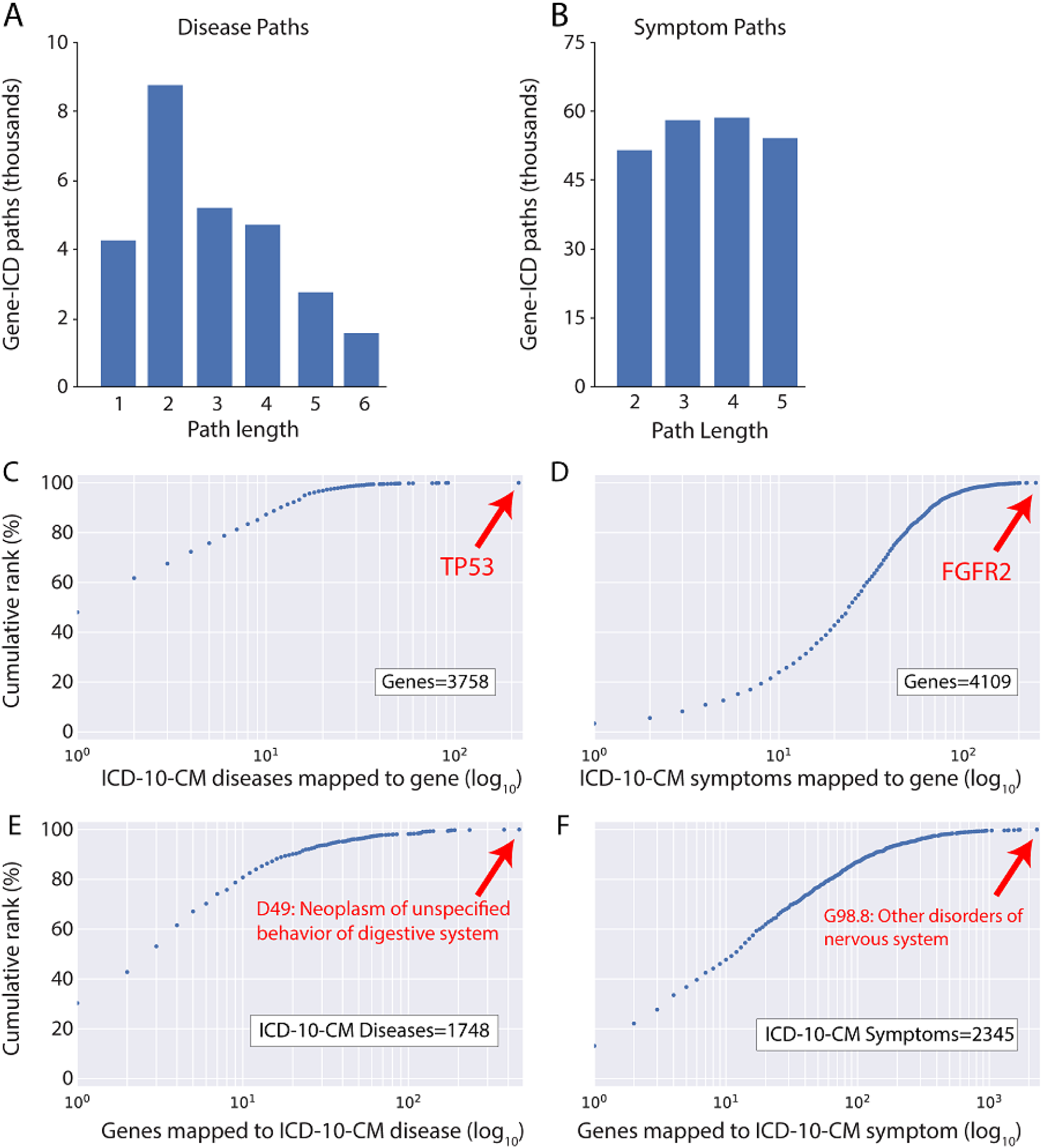
Results of unique gene-ICD paths for diseases and symptoms. A) Distribution of lengths for disease paths. B) Distribution of lengths for symptom paths. C-D) Cumulative rank distribution for counts of unique ICD-10-CM codes that map to each gene with C) disease paths and D) symptom paths. E-F) Cumulative rank distribution for counts of unique genes that map to each ICD-10-CM code with E) disease paths and F) symptom paths. Only paths to ICD-10-CM codes were included in C-F to prevent double counting of diagnoses. ICD codes are truncated to 4 digits and only 1 path for every gene-ICD pair is counted.

## 4. APPLICATION

To demonstrate the utility of our method, we applied our set of gene-ICD relationships to a cohort of 92,455 sequenced exomes paired with electronic health records (EHR) from the DiscovEHR collaboration[1]. Participants in the study contributed blood samples for exome sequencing and de-identified electronic health records according to the GHS Institutional Review Board–approved protocol (study number 2006-0258). Our goal was to use our mappings to validate the relationships between instances of predicted pathogenic variants in genes that are associated with disease and reports of diseases in the associated EHR. The EHR in this application consist of ICD-9-CM and ICD-10-CM codes listed at any time in the individual’s EHR (median record length of 14 years). We defined a predicted pathogenic variant as any potential loss-of-function (pLOF) mutation (stop loss, stop gain, start loss, frameshift, splice acceptor, splice donor) or a missense mutation that was predicted pathogenic by 5 variant pathogenicity prediction algorithms (SIFT[19], Polyphen2 HDIV and HVAR[20], LRT[21] and MutationTaster2[22]). Additionally, variants were filtered to those with a “pathogenic” or “likely pathogenic” clinical significance in ClinVar and had an allele frequency of 1% or less. Previous work using Geisinger MyCode data[1] used a broader set of criteria for selecting pathogenic variants and reported a median of 21 pLOFs per exome. We used a narrower set of criteria to enrich for variants with an expected pathogenic phenotype in their EHR, and this resulted in a median of 2 predicted pathogenic variants per exome. Additionally, we only used ICD codes that mapped to 20 or fewer genes with “disease” paths and 200 or fewer genes with “symptom” paths (this excluded ∼10% of ICD code mappings) to prevent excessive matches from ICD codes that map to a high number of genes. Using these criteria, the cohort contained 2,309 genes where at least one sample had one or more predicted pathogenic variants. Of these genes, our method was able to map 1,992 to at least one ICD code (Figure 5A). From these 1,992 genes, the cohort had 162,443 instances of a person-gene pair that had at least one variant (Figure 5B, outer circle), consisting of 10,355 unique variants. Using our mappings, we found valid associations between 816 genes (from 116 unique genes; 828 unique variants; 808 samples) and a “disease” ICD phenotype in the associated medical records (Figure 5B, left inner circle).

**Figure 5.**
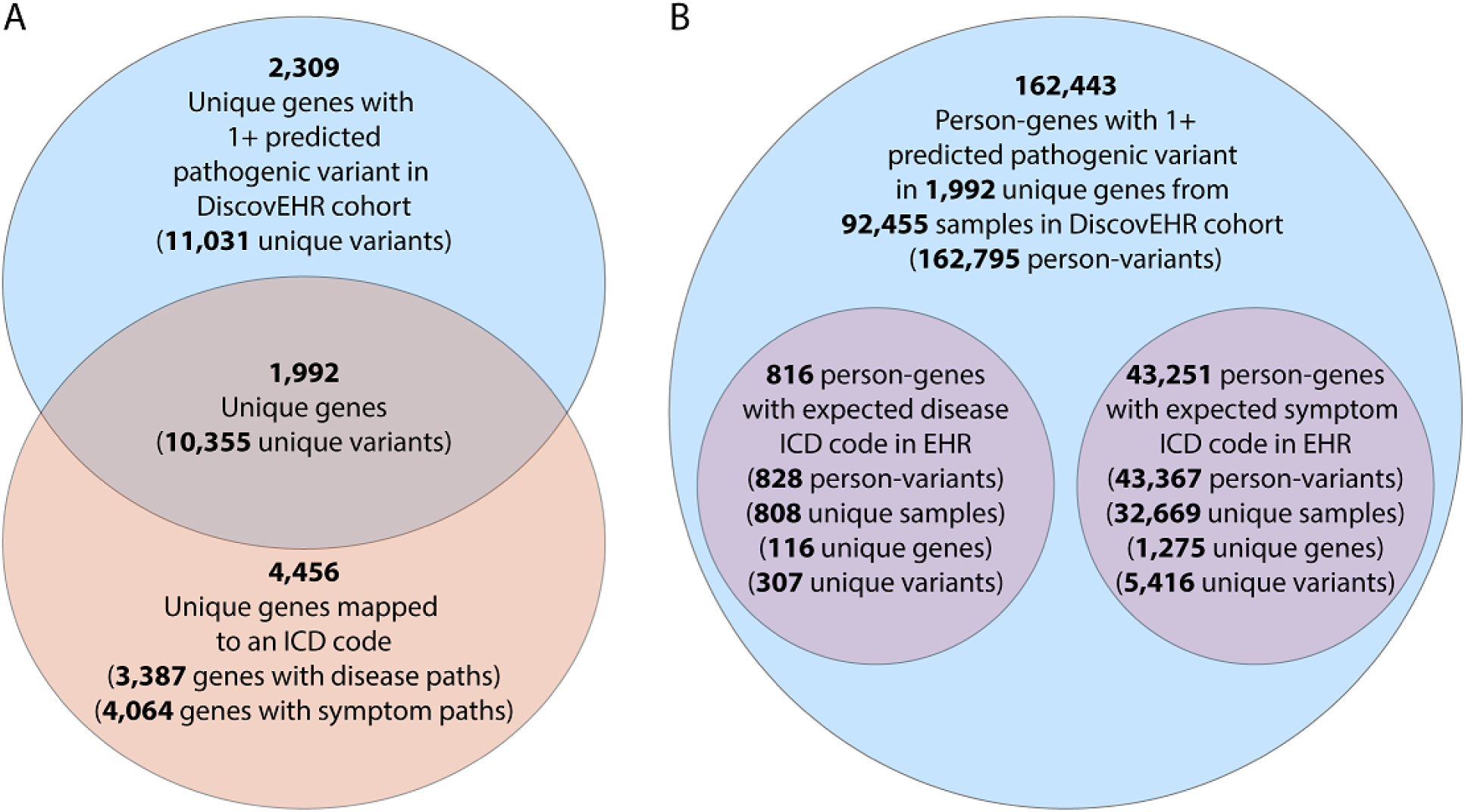
Euler diagrams showing pathogenic variants and associated diagnoses in a large cohort of sequenced exomes paired with electronic health records. A) Intersection of genes with pathogenic variants in DiscovEHR cohort and genes we have mapped to ICD codes. B) Outer circle is the total number of person-genes in the cohort with a pathogenic variant. Left inner circle is the number of person-genes with a pathogenic variant and the mapped “disease” ICD code in the individual’s electronic health records. Right inner circle is the number of person-genes with a pathogenic variant and the mapped “symptom” ICD code in the individual’s electronic health records. ICD code mappings include ICD-9-CM and ICD-10-CM. “Disease” and “symptom” paths where the ICD code maps to more than 20 or 200 genes, respectively, were excluded. Circles are not to scale.

There are several possible reasons why predicted pathogenic variants did not associate with expected ICD codes in the patients EHR, including[23, 24]:

- The variant could be due to an error in sequencing or variant calling
- Some benign variants in genes linked to disease have been incorrectly categorized as pathogenic
- Our gene-ICD mappings include both dominant and recessive conditions, and individuals with just one copy of a recessive pathogenic variant are not expected to develop the condition
- The individual’s genetic or environmental background can modify the effect of a variant
- Not all diseases will be diagnosed and documented in the patient’s EHR
- The disease could be captured in the clinical notes of the EHR, but not assigned an ICD code during processing
- The disease is present in the individual and captured in the EHR, but the ICD code selected by the physician was not mapped to the gene by our methodology

Additionally, we identified 43,251 of these genes (consisting of 1,275 unique genes; 43,367 variants; 32,669 samples) that also had an expected ICD “symptom” phenotype in their medical records (Figure 5B, right inner circle). Potential explanations for the difference between “disease” and “symptom” mappings include:

- Partial penetrance of recessive pathogenic variants can cause some symptoms but not the expected disease
- The disease was not diagnosed despite the presence and diagnosis of some symptoms
- Expected symptoms coinciding with an associated pathogenic variant by chance

In total, we found 32,786 of 92,455 samples in the cohort to have at least one “symptom” or “diagnosis” ICD code in their EHR that is mapped to a gene with a predicted pathogenic variant in their exome. Despite its limitations, this technique allowed us to rapidly identify individuals with a disease or symptom that is potentially caused by a genetic variant and flag the individual for further screening.

## 5. DISCUSSION

Our approach uses both the knowledge and semantic relationships embedded in biomedical vocabularies to validate clinically suspected genotype-phenotype relationships. Unlike other approaches to determining gene-disease relationships using the UMLS, we chose not to use Gene Ontology (GO) terms[25]. We made this choice not only because we are using our approach as a validation rather than a discovery tool, but also because we are interested in diseases (ICD codes) rather than specific genetic functions (GO terms). In future iterations of our approach, integrating GO or other function-based ontologies could help understand specific pathways or physiology involved in the gene-disease relationships it identifies. Additionally, because our approach is relatively generic in terms of starting and endpoints, one can employ it to examine relationships beyond genes and diseases. For example, we could use the set of relationships in the UMLS to map between genes and laboratory tests (different endpoints) or from the GWAS catalog or other disease vocabularies to specific disease-related ICD codes (different starting points).

Because of the breadth of the UMLS and the range of available relationships among concepts, and our goal of validating clinical gene-disease relationships, we used a specific set of filtering criteria and set relatively narrow limits on the gene-disease paths we accepted. Changing the stringency of the criteria could be appropriate depending on the use case. For example, looser criteria would be appropriate for discovery, while even more stringent criteria could be used if we were focused on a very narrow disease set or were trying to distinguish the true variant-disease relationships among a set of similar diseases. The ability to adjust the stringency of the criteria strengthens our approach as a method for identifying rare conditions that may have very specific clinical manifestations, particularly those that combine a set of specific symptoms and disease-related findings. The limitations related to validating rare diseases using EHR data are similar for any set of observational data, since ICD codes may be recorded with insufficient granularity and symptoms or other relevant findings are more often captured in clinical notes rather than in structured data.

There are several limitations to our approach. Most importantly, it will not create an exhaustive set of mappings between genes and diagnosis codes. The curation of the UMLS is semi-automated, and there is significant variation how thoroughly diseases are mapped to genes and how concepts are unified between sources. The result is that the number of phenotypes mapped to a gene is dependent on how thoroughly the gene is covered in the UMLS and its sources. Secondly, our method of validation involves a manual review to identify false mappings between gene and ICD code. However, the identification of missing mappings is far more challenging, and would likely require extensive input from experts on each disease.

Another validation approach would be to compare our results against a database with similar scope and aims; however, we are not aware of any that would offer a valid comparison. Many existing gene-phenotype resources are already incorporated into the UMLS or were incorporated into our database separately. This makes them a poor choice for comparison because each individual resource would be at best a subset of our results, and would be at an obvious disadvantage if they were compared directly. Additionally, many of these resources attempt to map diseases to a single “best” ICD phenotype (e.g., OMIM, Orphanet), or cover far fewer genes in more detail. A final caution about our method is that this it is not intended to substitute for human judgement. While we believe it is an effective way to highlight gene-phenotype relationships when the number of possible relationships is high, expert knowledge is still required for the interpretation of the relationship in the context of an individual’s genetics and medical records.

Our use of the UMLS to validate gene-disease relationships highlights some potential additions that would increase its usefulness as a tool from both the research and clinical perspective. For example, we found ∼600 concepts that indicated “susceptibility to” a disease but did not map to that specific disease. Additionally, we used MedGen and Orphanet to complement existing information in the UMLS specifically related to clinical manifestations of genetic diseases, but by using outside vocabularies alone we were unable to leverage all the concept relationships that exist in the UMLS. Finally, in developing and validating our paths through the UMLS, we found concept redundancies and missing concept mappings. Many of these originate with the source vocabularies; however, having the concepts as part of the UMLS now presents the opportunity to correct or supplement the mappings.

## 6. CONCLUSIONS

Though the UMLS links multiple biomedical vocabularies, its utility is often seen in mapping among clinical concepts generally in the research setting. By leveraging both the clinical and genetic knowledge embedded in the UMLS, we have shown its usefulness as a tool for validating gene-disease relationships found in a clinical setting. As more healthcare systems integrate clinical and genetic data, the demand for validating gene-disease relationships should only increase. Integrating additional vocabularies (e.g. MedGen and Orphanet) and validating and expanding existing gene-disease relationships already captured in the UMLS will help increase its utility as both a discovery and validation tool.

## ACKNOWLEDGEMENTS

We would like to thank the participants of the DiscovEHR Geisinger-Regeneron collaboration who have contributed biological samples and clinical records for research purposes. We would also like the thank Olivier Bodenreider and Kin Wah Fung for guidance on using the UMLS and the National Center for Biotechnology Information for creating the UMLS and MedGen and making them available as public resources.

## COMPETING INTERESTS

All individual authors are full-time employees of the Regeneron Genetics Center and receive stock in Regeneron Pharmaceuticals, Inc. as part of compensation. This research was funded by the Regeneron Genetics Center. No other conflicts are reported.

## AUTHOR CONTRIBUTIONS

A.L.B. developed the database, performed data analysis, created figures, wrote and revised manuscript. J.S. contributed figures, contributed to study design, wrote and revised manuscript. M.N.C contributed to study design, wrote and revised manuscript. C.G.-J., D.S., S.K. and S.B. contributed to study design, A.Y. contributed to preparation of phenotype data, J.G.R. and L.H. contributed bioinformatic resources.

## PRIOR PRESENTATION

Parts of this manuscript was presented as a poster at the American Society of Human Genetics Annual Meeting 2019 (Houston, Texas; October 15^th^-19^th^ 2019).

