## Supplemental tables and figures for "Validating gene-phenotype associations using relationships in the UMLS"

| **Source** | **File** | **Description** | **Rows** |
| --- | --- | --- | --- |
| UMLS | MRCONSO.RRF | A table of medical concepts, descriptions, and CUIs | 10,118,695 |
| MedGen | MGCONSO.RRF |  | 768,356 |
| UMLS | MRREL.RRF | A table of relationships between CUIs | 79,136,421 |
| MedGen | MGREL.RRF |  | 1,365,247 |
| UMLS | MRSTY.RRF | A table of semantic types for each CUI | 4,122,322 |
| MedGen | MGSTY.RRF |  | 427,247 |
| Orphanet | ORDO.csv | A table of rare diseases and their associated genes and ICD codes | 14,063 |

#### Table S1: Data sources to be included

| Data description | Source file | Data filtering | Column selection |
| --- | --- | --- | --- |
| All HGNC Gene names | MRCONSO.RRF | SUPPRESS=/=“O”  SAB=“HGNC”  TTY=“PT” | CODE, STR |
| All ICD-9-CM codes | MRCONSO.RRF / MGCONSO.RRF | SUPPRESS=/=“O”  SAB=“ICD9CM”  TTY=“PT” **OR** “HT” | CODE, STR |
| All ICD-10-CM codes | MRCONSO.RRF / MGCONSO.RRF | SUPPRESS=/=“O”  SAB=“ICD10CM”  TTY=“PT” **OR** “HT” | CODE, STR |
| Medical CUIs | MRCONSO.RRF / MGCONSO.RRF | SUPPRESS=/=“O”  SAB=Table S5 disease vocabulary abbreviations  TS=“P”  STT=“PF"  Filter CUIs by semantic type | CUI, STR |
| Medical CUIs semantic types | MRSTY.RRF / MGSTY.RRF | TUI=See table S3 for relevant semantic types | CUI, STY |

#### Table S2: Extracting the node data from UMLS sources. CODE = The HGNC code; STR = The HGNC gene name; CUI =UMLS CUI for the concept; STY = The semantic type

| Type of node | Semantic type code | Semantic type description |
| --- | --- | --- |
| Gene concepts | T028 | Gene or Genome |
|  | T049 | Cell or Molecular Dysfunction |
| Disease concepts | T019 | Congenital Abnormality |
|  | T028 | Gene or Genome |
|  | T033 | Finding |
|  | T046 | Pathologic Function |
|  | T047 | Disease or Syndrome |
|  | T048 | Mental or Behavioral Dysfunction |
|  | T049 | Cell or Molecular Dysfunction |
|  | T184 | Sign or Symptom |
|  | T190 | Anatomical Abnormality |
|  | T191 | Neoplastic Process |

#### Table S3: Semantic types for UMLS / MedGen concepts

| **Vocabulary** | **CUI** |
| --- | --- |
| Mesh | C0012674 |
| SNOMEDCT_US | C0037088 |
| HPO | C4021819 |
| CCS10 | C0011900 |
| NCI | C1511989 |
| MEDCIN | C0012634 |
| MEDRA | C1140263 |
| ICD9CM | Any CUI that maps to an ICD-9-CM code range |
| ICD10CM | Any CUI that maps to an ICD-10-CM code range |

#### Table S4: Header concepts for each vocabulary

| Data description | Source file | Data filtering | Column selection | Subtypes |
| --- | --- | --- | --- | --- |
| HGNC Genes to CUIs | MRCONSO.RRF / MGCONSO.RRF | SUPPRESS=/=“O”  SAB=“HGNC”  TTY=“PT” | CODE, CUI | No |
| CUIs to ICD-9-CM codes | MRCONSO.RRF / MGCONSO.RRF | SUPPRESS=/=“O”  SAB=“ICD9CM”  TTY=“PT” **OR** “HT” | CODE, CUI | No |
| CUIs to ICD-10-CM codes | MRCONSO.RRF / MGCONSO.RRF | SUPPRESS=/=“O”  SAB=“ICD10CM”  TTY=“PT” **OR** “HT” | CODE, CUI | No |
| Edges between CUIs | MRREL.RRF / MGREL.RRF | SUPPRESS=/=“O”  SAB=Table S6 disease vocabulary abbreviations | CUI1, CUI2, SAB, REL, RELA | Yes, see tables S6, S7 |

#### Table S5: Edge data extracted from UMLS sources. CODE = The HGNC or ICD code; CUI = The CUI code; CUI1 = Source CUI code; CUI2 = Destination CUI code; SAB = The source vocabulary; REL = The relationship; RELA = The relationship attribute

| Type of node | Vocabulary abbreviation | Vocabulary description |
| --- | --- | --- |
| Gene concepts | HGNC | HUGO Gene Nomenclature Committee |
| UMLS concepts | CCS_10 | Clinical Classifications Software 10 |
|  | GTR | Genetic Testing Registry |
|  | HGNC | HUGO Gene Nomenclature Committee |
|  | HPO | Human Phenotype Ontology |
|  | ICD10CM | International Classification of Diseases, Tenth Revision, Clinical Modification |
|  | ICD9CM | International Classification of Diseases, Ninth Revision, Clinical Modification |
|  | MDR | Medical Dictionary for Regulatory Activities |
|  | MEDCIN | MEDCIN |
|  | MSH | Medical Subject Headings |
|  | MTH | Metathesaurus Names |
|  | NCI | National Cancer Institute Thesaurus |
|  | OMIM | Online Mendelian Inheritance in Man |
|  | SNOMEDCT_US | SNOMED CT, US Edition |
| ICD Concepts | ICD10CM | International Classification of Diseases, Tenth Revision, Clinical Modification |
|  | ICD9CM | International Classification of Diseases, Ninth Revision, Clinical Modification |

#### Table S6: Ontology and vocabulary concept filters for UMLS / MedGen concept relationships

| Source | Relationship | Relationship attribute | Filter code |
| --- | --- | --- | --- |
| CCS_10 | CHD | null | B |
| CCS_10 | PAR | null | G |
| CCS_10 | RQ | classified_as | B |
| CCS_10 | RQ | classifies | G |
| CCS_10 | SIB | null | E |
| CCS_10 | SY | has_multi_level_category | Syn |
| CCS_10 | SY | has_single_level_category | Syn |
| GTR | CHD | null | B |
| GTR | PAR | null | G |
| GTR | RO | related_to | E |
| HPO | CHD | isa | B |
| HPO | CHD | null | B |
| HPO | PAR | inverse_isa | G |
| HPO | PAR | null | G |
| HPO | RB | null | G |
| HPO | RN | null | B |
| HPO | RO | clinical_course_of | E |
| HPO | RO | has_clinical_course | E |
| HPO | RO | has_inheritance_type | E |
| HPO | RO | has_manifestation | E |
| HPO | RO | inheritance_type_of | E |
| HPO | RO | manifestation_of | Sym |
| HPO | RO | null | U |
| HPO | RQ | consider | E |
| HPO | RQ | consider_from | Syn |
| HPO | RQ | replaced_by | E |
| HPO | RQ | replaces | Syn |
| HPO | SIB | null | E |
| HPO | SY | null | Syn |
| ICD10CM | CHD | null | B |
| ICD10CM | PAR | null | G |
| ICD10CM | RQ | null | E |
| ICD10CM | SIB | null | E |
| ICD9CM | CHD | null | B |
| ICD9CM | PAR | null | G |
| ICD9CM | SIB | null | E |
| MDR | CHD | null | B |
| MDR | PAR | null | G |
| MDR | RQ | classified_as | E |
| MDR | RQ | classifies | G |
| MDR | SIB | null | E |
| MEDCIN | CHD | isa | B |
| MEDCIN | PAR | inverse_isa | G |
| MEDCIN | RB | inverse_isa | G |
| MEDCIN | RN | isa | B |
| MEDCIN | RO | associated_finding_of | E |
| MEDCIN | RO | associated_morphology_of | G |
| MEDCIN | RO | associated_with | E |
| MEDCIN | RO | cause_of | G |
| MEDCIN | RO | component_of | G |
| MEDCIN | RO | definitional_manifestation_of | G |
| MEDCIN | RO | direct_morphology_of | G |
| MEDCIN | RO | due_to | Sym |
| MEDCIN | RO | finding_context_of | E |
| MEDCIN | RO | finding_site_of | E |
| MEDCIN | RO | focus_of | G |
| MEDCIN | RO | has_associated_finding | E |
| MEDCIN | RO | has_associated_morphology | E |
| MEDCIN | RO | has_component | E |
| MEDCIN | RO | has_definitional_manifestation | E |
| MEDCIN | RO | has_direct_morphology | E |
| MEDCIN | RO | has_finding_context | E |
| MEDCIN | RO | has_finding_site | E |
| MEDCIN | RO | has_focus | E |
| MEDCIN | RO | has_indirect_procedure_site | E |
| MEDCIN | RO | has_interpretation | E |
| MEDCIN | RO | has_pathological_process | E |
| MEDCIN | RO | has_procedure_context | E |
| MEDCIN | RO | has_procedure_morphology | E |
| MEDCIN | RO | has_severity | E |
| MEDCIN | RO | indirect_procedure_site_of | E |
| MEDCIN | RO | interpretation_of | E |
| MEDCIN | RO | interprets | E |
| MEDCIN | RO | is_interpreted_by | E |
| MEDCIN | RO | occurs_after | E |
| MEDCIN | RO | occurs_before | G |
| MEDCIN | RO | pathological_process_of | G |
| MEDCIN | RO | procedure_context_of | E |
| MEDCIN | RO | procedure_morphology_of | E |
| MEDCIN | RO | severity_of | E |
| MEDCIN | SIB | null | E |
| MEDCIN | SY | same_as | Syn |
| MSH | AQ | null | E |
| MSH | CHD | null | B |
| MSH | PAR | null | G |
| MSH | QB | null | E |
| MSH | RB | mapped_from | G |
| MSH | RB | null | G |
| MSH | RN | mapped_to | B |
| MSH | RN | null | B |
| MSH | RO | has_mapping_qualifier | E |
| MSH | RO | mapping_qualifier_of | E |
| MSH | RO | null | U |
| MSH | SIB | null | E |
| MTH | RB | null | G |
| MTH | RN | null | B |
| MTH | RO | null | U |
| NCI | CHD | isa | B |
| NCI | PAR | inverse_isa | G |
| NCI | RB | mapped_from | G |
| NCI | RB | mapped_to | G |
| NCI | RN | mapped_from | B |
| NCI | RN | mapped_to | B |
| NCI | RO | abnormality_associated_with_allele | G |
| NCI | RO | allele_has_abnormality | E |
| NCI | RO | allele_in_chromosomal_location | E |
| NCI | RO | biological_process_has_initiator_process | E |
| NCI | RO | biological_process_has_result_biological_process | E |
| NCI | RO | biological_process_is_part_of_process | E |
| NCI | RO | biological_process_results_from_biological_process | G |
| NCI | RO | cdrh_parent_of | G |
| NCI | RO | chromosomal_location_of_allele | E |
| NCI | RO | concept_in_subset | E |
| NCI | RO | ctcae_5_parent_of | G |
| NCI | RO | disease_excludes_cytogenetic_abnormality | E |
| NCI | RO | disease_excludes_finding | E |
| NCI | RO | disease_excludes_molecular_abnormality | E |
| NCI | RO | disease_has_abnormal_cell | E |
| NCI | RO | disease_has_associated_disease | E |
| NCI | RO | disease_has_associated_gene | G |
| NCI | RO | disease_has_cytogenetic_abnormality | E |
| NCI | RO | disease_has_finding | E |
| NCI | RO | disease_has_molecular_abnormality | G |
| NCI | RO | disease_is_grade | E |
| NCI | RO | disease_is_marked_by_gene | G |
| NCI | RO | disease_is_stage | E |
| NCI | RO | disease_mapped_to_gene | G |
| NCI | RO | disease_may_have_abnormal_cell | E |
| NCI | RO | disease_may_have_associated_disease | E |
| NCI | RO | disease_may_have_cytogenetic_abnormality | G |
| NCI | RO | disease_may_have_finding | E |
| NCI | RO | disease_may_have_molecular_abnormality | G |
| NCI | RO | eo_disease_has_property_or_attribute | B |
| NCI | RO | eo_disease_maps_to_human_disease | B |
| NCI | RO | gene_associated_with_disease | B |
| NCI | RO | gene_encodes_gene_product | B |
| NCI | RO | gene_has_abnormality | B |
| NCI | RO | gene_in_chromosomal_location | B |
| NCI | RO | gene_involved_in_molecular_abnormality | B |
| NCI | RO | gene_involved_in_pathogenesis_of_disease | B |
| NCI | RO | gene_is_biomarker_of | B |
| NCI | RO | gene_is_element_in_pathway | B |
| NCI | RO | gene_mapped_to_disease | B |
| NCI | RO | gene_mutant_encodes_gene_product_sequence_variation | B |
| NCI | RO | gene_plays_role_in_process | B |
| NCI | RO | gene_product_encoded_by_gene | B |
| NCI | RO | gene_product_sequence_variation_encoded_by_gene_mutant | B |
| NCI | RO | genetic_biomarker_related_to | B |
| NCI | RO | has_cdrh_parent | B |
| NCI | RO | has_ctcae_5_parent | B |
| NCI | RO | has_inc_parent | B |
| NCI | RO | has_nichd_parent | B |
| NCI | RO | human_disease_maps_to_eo_disease | B |
| NCI | RO | inc_parent_of | G |
| NCI | RO | is_abnormal_cell_of_disease | G |
| NCI | RO | is_abnormality_of_gene | G |
| NCI | RO | is_associated_disease_of | E |
| NCI | RO | is_chromosomal_location_of_gene | E |
| NCI | RO | is_cytogenetic_abnormality_of_disease | G |
| NCI | RO | is_finding_of_disease | Sym |
| NCI | RO | is_grade_of_disease | E |
| NCI | RO | is_molecular_abnormality_of_disease | E |
| NCI | RO | is_not_cytogenetic_abnormality_of_disease | E |
| NCI | RO | is_not_finding_of_disease | E |
| NCI | RO | is_not_molecular_abnormality_of_disease | E |
| NCI | RO | is_property_or_attribute_of_eo_disease | E |
| NCI | RO | is_stage_of_disease | E |
| NCI | RO | mapped_from | E |
| NCI | RO | mapped_to | G |
| NCI | RO | may_be_abnormal_cell_of_disease | Sym |
| NCI | RO | may_be_associated_disease_of_disease | Sym |
| NCI | RO | may_be_cytogenetic_abnormality_of_disease | E |
| NCI | RO | may_be_finding_of_disease | Sym |
| NCI | RO | may_be_molecular_abnormality_of_disease | E |
| NCI | RO | molecular_abnormality_involves_gene | G |
| NCI | RO | neoplasm_has_special_category | E |
| NCI | RO | nichd_parent_of | G |
| NCI | RO | pathogenesis_of_disease_involves_gene | G |
| NCI | RO | pathway_has_gene_element | G |
| NCI | RO | process_includes_biological_process | G |
| NCI | RO | process_initiates_biological_process | E |
| NCI | RO | process_involves_gene | G |
| NCI | RO | related_to_genetic_biomarker | G |
| NCI | RO | special_category_includes_neoplasm | G |
| NCI | RO | subset_includes_concept | G |
| NCI | SY | mapped_from | Syn |
| NCI | SY | mapped_to | Syn |
| OMIM | CHD | null | B |
| OMIM | PAR | null | G |
| OMIM | RO | has_inheritance_type | E |
| OMIM | RO | has_manifestation | E |
| OMIM | RO | has_phenotype | B |
| OMIM | RO | inheritance_type_of | E |
| OMIM | RO | manifestation_of | Sym |
| OMIM | RO | phenotype_of | G |
| OMIM | RQ | alias_of | Syn |
| OMIM | RQ | allelic_variant_of | G |
| OMIM | RQ | has_alias | Syn |
| OMIM | RQ | has_allelic_variant | E |
| ORDO | CHD | part_of | B |
| ORDO | PAR | has_part | G |
| ORDO | RO | has_inheritance_type | E |
| ORDO | RO | inheritance_type_of | E |
| SNOMEDCT_US | CHD | isa | B |
| SNOMEDCT_US | PAR | inverse_isa | G |
| SNOMEDCT_US | RB | alternative_of | G |
| SNOMEDCT_US | RB | inverse_was_a | G |
| SNOMEDCT_US | RB | referred_to_by | G |
| SNOMEDCT_US | RN | has_alternative | B |
| SNOMEDCT_US | RN | refers_to | B |
| SNOMEDCT_US | RN | was_a | B |
| SNOMEDCT_US | RO | associated_etiologic_finding_of | E |
| SNOMEDCT_US | RO | associated_finding_of | E |
| SNOMEDCT_US | RO | associated_function_of | E |
| SNOMEDCT_US | RO | associated_morphology_of | E |
| SNOMEDCT_US | RO | associated_procedure_of | E |
| SNOMEDCT_US | RO | associated_with | E |
| SNOMEDCT_US | RO | causative_agent_of | E |
| SNOMEDCT_US | RO | cause_of | G |
| SNOMEDCT_US | RO | clinical_course_of | E |
| SNOMEDCT_US | RO | course_of | E |
| SNOMEDCT_US | RO | definitional_manifestation_of | G |
| SNOMEDCT_US | RO | direct_morphology_of | E |
| SNOMEDCT_US | RO | due_to | Sym |
| SNOMEDCT_US | RO | entire_anatomy_structure_of | E |
| SNOMEDCT_US | RO | finding_context_of | E |
| SNOMEDCT_US | RO | finding_site_of | E |
| SNOMEDCT_US | RO | focus_of | G |
| SNOMEDCT_US | RO | has_associated_etiologic_finding | E |
| SNOMEDCT_US | RO | has_associated_finding | E |
| SNOMEDCT_US | RO | has_associated_function | E |
| SNOMEDCT_US | RO | has_associated_morphology | E |
| SNOMEDCT_US | RO | has_associated_procedure | E |
| SNOMEDCT_US | RO | has_causative_agent | E |
| SNOMEDCT_US | RO | has_clinical_course | E |
| SNOMEDCT_US | RO | has_course | E |
| SNOMEDCT_US | RO | has_definitional_manifestation | E |
| SNOMEDCT_US | RO | has_direct_morphology | E |
| SNOMEDCT_US | RO | has_entire_anatomy_structure | G |
| SNOMEDCT_US | RO | has_finding_context | E |
| SNOMEDCT_US | RO | has_finding_site | E |
| SNOMEDCT_US | RO | has_focus | E |
| SNOMEDCT_US | RO | has_interpretation | E |
| SNOMEDCT_US | RO | has_pathological_process | E |
| SNOMEDCT_US | RO | has_precondition | E |
| SNOMEDCT_US | RO | has_procedure_context | E |
| SNOMEDCT_US | RO | has_realization | E |
| SNOMEDCT_US | RO | has_severity | E |
| SNOMEDCT_US | RO | interpretation_of | E |
| SNOMEDCT_US | RO | interprets | E |
| SNOMEDCT_US | RO | is_interpreted_by | E |
| SNOMEDCT_US | RO | mapped_from | E |
| SNOMEDCT_US | RO | mapped_to | E |
| SNOMEDCT_US | RO | occurs_after | Sym |
| SNOMEDCT_US | RO | occurs_before | G |
| SNOMEDCT_US | RO | pathological_process_of | G |
| SNOMEDCT_US | RO | plays_role | E |
| SNOMEDCT_US | RO | possibly_equivalent_to | E |
| SNOMEDCT_US | RO | precondition_of | E |
| SNOMEDCT_US | RO | procedure_context_of | E |
| SNOMEDCT_US | RO | realization_of | G |
| SNOMEDCT_US | RO | replaced_by | E |
| SNOMEDCT_US | RO | replaces | Syn |
| SNOMEDCT_US | RO | role_played_by | E |
| SNOMEDCT_US | RO | severity_of | E |
| SNOMEDCT_US | RO | temporally_followed_by | E |
| SNOMEDCT_US | RO | temporally_follows | E |
| SNOMEDCT_US | SIB | null | E |
| SNOMEDCT_US | SY | same_as | Syn |

#### Table S7: Filtering of relationship and relationship attribute code pairs for inclusion in database

#### Note S1: Exception for edge filtering

Table S7 lists the prioritization scheme for edges based on the source ontology, relationship and relationship attribute. One exception to this filtering is a set of edges from the NCI ontology. While the majority of the edges in that are correct, there are several thousand gene variant nodes that should have their filter code set to “G”. They can be identified by:

1) SAB=”NCI”
2) REL=”CHD”
3) RELA=”isa”
4) Semantic type is “Gene or Genome”
5) Map uniquely to a CUI of a gene name

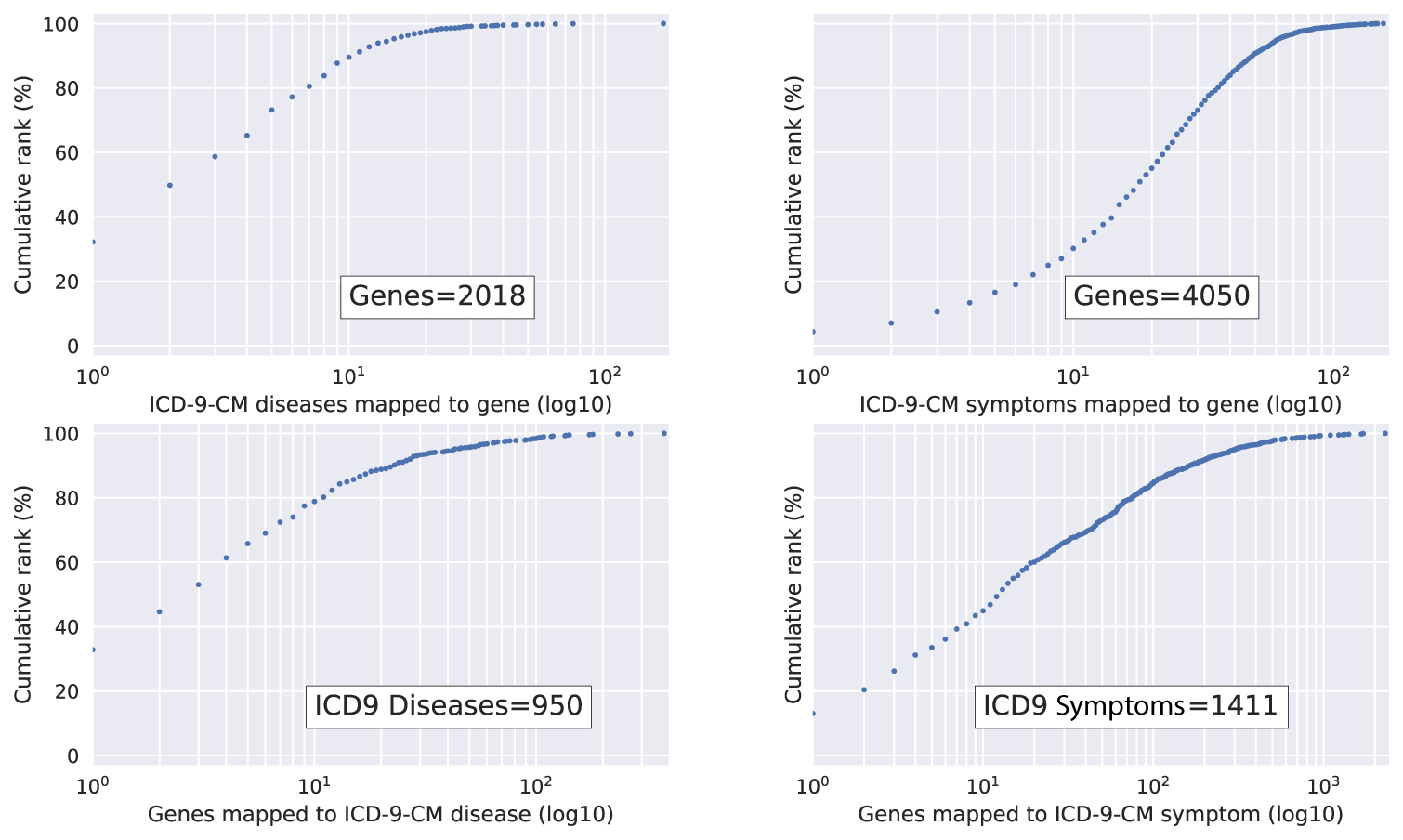

#### Figure S1. Specificity of gene and ICD mappings for ICD-9-CM codes

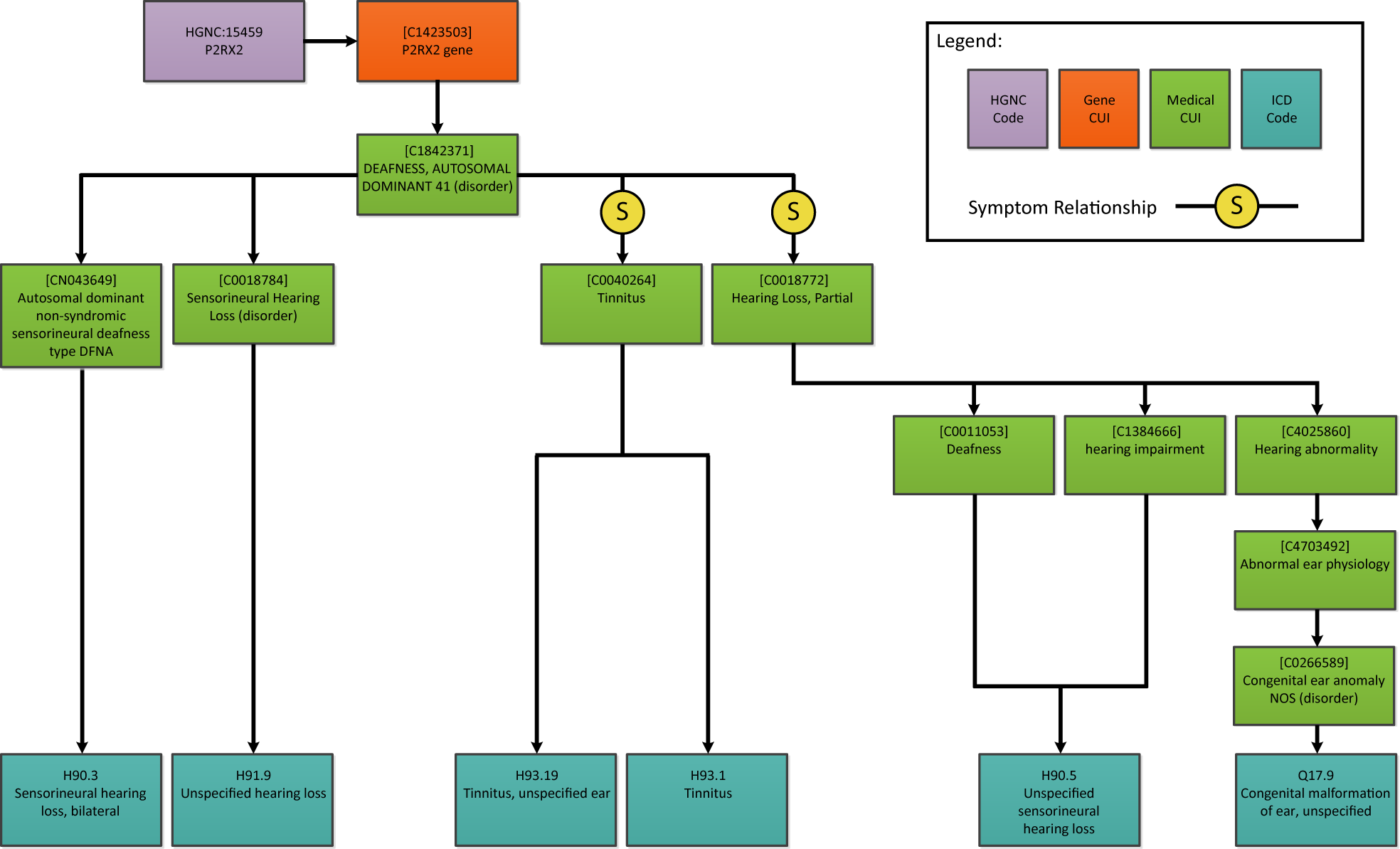

#### Figure S2: ICD Diagnosis codes mappings for gene P2RX2
